# Mice Models to Bioinformatics Methods Led to the Discovery of Antidiabetic Compounds in the *F. racemosa* Plant Extracts

**DOI:** 10.1101/2023.11.22.568234

**Authors:** Mohammad Uzzal Hossain, ABZ Naimur Rahman, Md. Shahadat Hossain, Shajib Dey, Zeshan Mahmud Chowdhury, Arittra Bhattacharjee, Ishtiaqe Ahmed, Mohammad Kamrul Hasan, Istiak Ahmed, Md. Billal Hosen, Keshob Chandra Das, Chaman Ara Keya, Md. Salimullah

## Abstract

One of the primary health issues caused by inadequate blood sugar regulation is Diabetes Mellitus (DM). Diabetes and its consequences remain clinically significant even with the development of oral hypoglycemic medications. Therefore, *Ficus racemosa* (*F. racemosa*) plant has been studied for assessing of its antidiabetic potential coupling with animal model and *in silico* experiments. Drug Alloxan (150 mg/kg) was injected to induce the experimental diabetes in Swiss Albino mice, and two doses methanol extract of the *F. racemosa* fruit (300 and 500 mg/kg) along with glibenclamide (5 mg/kg) were given orally. Oral Glucose Tolerance Test (OGTT) and acute toxicity were performed as well in both diabetic and non-diabetic mice. Later, *in silico* experiments including ADMET profiling, molecular docking and simulations were performed. The administration of a dosage less than 3000 mg/kg has been observed to be well-tolerated by mice, with no reported instances of mortality or adverse effects. Following oral administration for 7 days, the blood glucose level (BGL) was significantly decreased in mice model in both doses of extracts, indicating the effect of *F. racemosa*. Subsequent to this, molecular docking and simulations have indicated that the SIRT1 receptor exhibits a higher binding affinity towards four specific compounds, namely Friedelin, Lupeol Acetate, Gluanol, and Ferulic Acid, as indicated by the dynamics parameters and interacting residues. The current investigation provided evidence that the fruit extract of *F. racemosa* significantly mitigated the hyperglycemic impact. Moreover, a total of four substances have been found that play a crucial role in the mechanisms behind the reduction of diabetic effects. Hence, the current investigation could potentially serve as a viable therapeutic approach in the treatment of diabetes.

## 1. Introduction

Type 2 diabetes is a chronic medical condition characterized by high blood sugar levels due to the body’s inability to use insulin effectively (1). It is typically associated with obesity, physical inactivity, and a diet high in processed foods and sugar (2). Over time, high blood sugar levels can damage organs such as the kidneys, eyes, and nerves (3). Type 2 diabetes can often be managed through lifestyle changes such as healthy eating, regular exercise, and weight loss (4). Medications such as metformin, sulfonylureas, and SGLT-2 inhibitors may also be prescribed to help control blood sugar levels (3,5). Regular monitoring of blood sugar levels and close communication with a healthcare provider are important to prevent or delay the onset of complications for both types of diabetes (4). Diabetes is a chronic disease that affects millions of people worldwide (6). According to the International Diabetes Federation (IDF), in 2021, there were an estimated 537 million adults living with diabetes globally. This number is expected to be 642 million by 2040 (7). The prevalence of diabetes varies by region, with the Western Pacific and Southeast Asian regions having the highest number of cases (8). Diabetes is a leading cause of death, with 4.2 million deaths attributed to the disease in 2019 (9). In addition to its impact on mortality, diabetes also imposes a significant economic burden. In 2019, the global cost of diabetes was estimated to be $760 billion USD (10). Despite the availability of various treatment, diabetes remains a significant health concern, affecting millions of people worldwide (American Diabetes Association, 2021; International Diabetes Federation, 2019) (1,11). Current treatments may have side effects and are not always effective in controlling blood sugar levels in all patients (12). Therefore, there is a need for alternative treatments that are safe, effective, and easily accessible to help manage and prevent complications associated with diabetes (13).

*Ficus racemosa*, commonly known as the cluster fig tree or Gular, is a large, deciduous tree species found in tropical and subtropical regions, including India, Pakistan, and Sri Lanka (14). The plant has a long history of use in Ayurvedic medicine for various purposes, including the treatment of diabetes (15). For ages, its leaves, bark, and fruit have been utilized in traditional medicine, with the plant thought to have anti-inflammatory, anti-diabetic, and anti-arthritic effects (14). The plant has also been used to treat skin diseases, respiratory problems, and gastrointestinal disorders (14). The chemical composition of the *F. racemosa* plant has been extensively studied, with researchers identifying a range of bioactive compounds, including flavonoids, alkaloids, terpenoids, and phenolic compounds (16). It is widely believed that these compounds have a significant role in the plant’s diverse medical properties, which encompass its potential as an agent for managing diabetes. Several studies have found that the plant’s extracts may help lower blood sugar levels and improve insulin sensitivity, potentially making it a beneficial therapeutic choice for diabetics (17). While further research is needed to fully understand the plant’s mechanisms of action and safety profile, these findings indicate that *F. racemosa. F. racemosa* may have promising potential as a natural antidiabetic agent.

Studies have also examined the potential of *F. racemosa* plant compounds as an alternative to insulin therapy in diabetes management. These compounds improve glucose metabolism, and insulin sensitivity, and promote glucose uptake (18). In comparison to insulin therapy, *F. racemosa* compounds may have fewer side effects, a broader therapeutic range, and a lower risk of hypoglycemia and weight gain (19). However, additional research is needed to comprehend their mechanisms of action, safety profile, and long-term efficacy. Despite the challenges, plant compounds derived from *F. racemosa* may provide a new avenue for managing diabetes, particularly in settings with limited resources (18). Previous studies on *F. racemosa* and diabetes have been limited by conventional methods, leaving a research gap in understanding the specific compounds and their effects. Our research used contemporary approaches, combining wet lab and dry lab techniques, to investigate the potential of *Ficus racemosa*. We found that the compound had a significant positive impact on insulin sensitivity and glucose uptake, highlighting its potential as a treatment for diabetes.

## 2. Method and materials

### 2.1 Collection and preparation of plant materials

In this study, fresh fruits of *F. racemosa* were collected from Rangamati, a district located within the Chittagong hill tracts of Bangladesh. The taxonomic identification and confirmation of the plant species were carried out with precision by the Plant Biotechnology Division at the National Institute of Biotechnology. The fruits were subjected to a series of thorough rinses with distilled water in order to remove any dust particles. After a week of air-drying in the shade at room temperature, the dried leaves were finely powdered.

The dry, powdered form of the fruits was then mixed properly and dissolved in a methanol solution. The flask was left at room temperature for 72 hours with a constant magnetic stirrer. Afterward, the mixture underwent filtration, and methanol was subsequently eliminated using a rotary evaporator, resulting in the production of the crude methanolic extract. For the experiment, the freeze-dried methanolic extract was preserved in the fridge. The dried plant crude extract was then reconstituted with an appropriate solvent for oral administration.

### 2.2 Experimental animals

Both adult female and male Swiss albino mice aged eight weeks old and weighing 24-35 g were obtained from the International Centre for Diarrhoeal Disease Research (icddr,b), Bangladesh. The mice were raised in polypropylene cages of the animal house. The house was maintained at a temperature of 23 ± 2°C with a 12h/12h light/dark cycle (20–23). The animals were acclimatized for 1 week before the experiment was carried out. They were provided with standard food pellets and water throughout the experiment apart from the fasting period. The study protocol obtained approval from the Plant Biotechnology Division at the National Institute of Biotechnology ensuring that the treatment and care of the animals adhered to ethical standards for laboratory animal use (23).

### 2.3 Acute toxicity test

Acute toxicity tests were conducted on the crude extract following an overnight fast, during which the animals had only access to water (24). The study consisted of five groups, with each group consisting of eight mice. The control group received only the vehicle while the treatment groups were administered 1000, 2000, and 3000 mg/kg of the *F. racemosa* crude extract (25). During the first four hours following extract administration, the animals were carefully monitored for any observable signs of toxicity or unusual behaviors such as agitation, tremors, diarrhea, lethargy, weight loss or paralysis (26). Following that, they were observed daily for two weeks to assess any changes in their overall behavior or physical activity. Food was supplied again four hours after the extract was administered.

### 2.4 Induction of experimental diabetes

A comprehensive assessment was conducted on Swiss albino mice after a period of fasting which lasted between 12 to 14 hours. After this, the mice were administered a single intraperitoneal injection of freshly prepared alloxan monohydrate solution (150 mg/kg body weight), which led to the development of diabetes in each of them. A digital weight balance was used to gauge their body weight, and a glucometer was used to assess their fasting blood sugar levels. The alloxan solution was precisely prepared for injection, involving a 0.5 mL mixture with pH 4.5 sodium citrate. Food and water were given to the animals thirty minutes after administering alloxan (27–30). After a 48-hour period post-alloxan injection, we meticulously measured the plasma blood glucose levels of each animal. This was done using blood droplets obtained from the tail using a sharp single-use needle. Only animals having a fasting blood glucose level above 200 mg/dL were included in the study (29,30).

### 2.5 Experimental design

The five groups of mice (both male and female) were subjected to the crude extract and standard control drug (Glibenclamide). One of the five groups was allocated as the control group. The extract was given to both male and female mice at a dose of 300mg/kgb.w t, and the effects were evaluated during a 4 hour period of treatment for 7 days. Normal control group (NC) is the normal control group, which is given 10 mg/kg of normal saline. Diabetic control (DC) is the diabetic control group that receives the vehicle. Glibenclamide control (GC) is the positive control group that receives glibenclamide medication at a dose of 5 mg/kg. FR300 and FR500 are the treatment groups that receive *F. racemosa* extract at a dose of 300 and 500 mg/kg.

### 2.6 Oral glucose tolerance test (OGTT)

Prior to conducting the oral glucose tolerance test (OGTT), the animals underwent a 24-hour period of fasting, during which they were supplied with unlimited access to water. Subsequently, a 1 mL/kg dosage of oral glucose solution (2 g/kg body weight) was given. Following glucose administration, blood samples were collected from each mouse at 30 minutes intervals (0, 30, 60 and 120 minutes (20).

### 2.7 Preparation of target protein and selection of plant compounds

The 3D crystal structure “4I5I” of the Homo sapiens target protein SIRT1 was obtained from the Protein Data Bank (PDB) maintained by the Research Collaboratory for Structural Bioinformatics (RCSB) PDB (31). To ensure that the data could be used for further studies, the PDB file format of SIRT1 *Homo sapiens* was preserved. Accordingly, all available literature and databases were searched for plant species with antidiabetic properties that were supported by laboratory research, either for the plant itself or its constituent compounds. Following an evaluation of various characteristics, a total of 13 different compounds were selected from the *F. racemosa* plant. The structure of the chosen active chemicals of this plant was obtained from the PubChem database in the SDF (Structure Data File) format as a three-dimensional (3D) structure. Additionally, PubChem was used to obtain the two-dimensional structure of the compounds (32).

### 2.8 Pharmacoinformatics elucidation (ADMET Profiling)

The pharmacokinetic properties like absorption, distribution, metabolism, excretion, (ADME) profiling of 13 active compounds of *F. racemosa* fruit were determined using the ADMETlab 2.0 and pkCSM web descriptive servers (33,34). The ADMETlab 2.0 were used to analyze the adsorption, metabolism, and excretion and the pkCSM was employed to evaluate the distribution properties of those compounds. The pkCSM utilizes graph-based signatures for the establishment of ADMET features and aid in drug development. The toxicity parameters like hepatotoxicity, carcinogenicity, immunotoxicity and mutagenicity of the 13 compounds were calculated using the ProTox-II server. (**Supplementary Figure**) ProTox-II server predicts various toxicity endpoints using the different machine-learning models, molecular similarities and pharmacophores.

### 2.9 Energy minimization of the protein

Since the drawn chemical structures are not energetically favorable, it is essential to determine the correct molecular arrangement in space through the process of energy minimization. To achieve this, the Swiss PDB Viewer (SPDBV) was used. SPDBV is a multi-platform protein structure visualization program that enables computation and visualization of the electrostatic potentials of proteins, as well as the capability to model protein structures (35). Minimization of the target protein, the energy minimization process of SIRT1 was carried out using SPDBV that demonstrated the minimized form of the SIRT1 target protein.

### 2.10 Structural optimization of the compounds

Structural optimization of the selected compounds was performed using Avogadro, a powerful open-source molecular modeling software (36). Avogadro provides a wide range of tools for structural optimization, including force field optimization, Monte Carlo simulations, and genetic algorithms. This tool was used to find the most stable and energetically favorable structures for the selected 13 active compounds.

### 2.11 Molecular docking

The molecular docking analysis of four selected compounds was carried out using PyRx tools Autodock vina to determine the most stable binding conformation (37). For each compound, the dimensions (X = 65, Y = 40, and Z = 50) and center (X = 20.527, Y = -57.458, and Z = 27.219) of the grid box were set, as well as the grid box spacing (1.0). Resultant docked compounds that have better binding affinity (kcal/mol) were retrieved and visualized by using pyMOL and BIOVA Discovery Studio Visualizer Tool (38,39).

### 2.12 Molecular Dynamics Simulation

Molecular dynamics simulations were performed for using GROningen MAchine for Chemical Simulations aka GROMACS (version 5.1.1) to study the dynamic interactions under specific physiological conditions between SIRT1 protein and SIRT1-Ferulic acid complex, SIRT1-Friedeline complex, SIRT1-Gluanol complex and SIRT1-Lupeol acetate complex (40). The radius of gyration (Rg), root mean square deviation (RMSD), root mean square fluctuation (RMSF) and solvent accessible surface area (SASA) data were calculated for assessing the binding stability, conformational changes, and key intermolecular interactions.

## 3. Results

The overall workflow has been demonstrated in **Fig 1** and the location of the sample collection has been shown in **Fig 2**.

**Figure 1:**
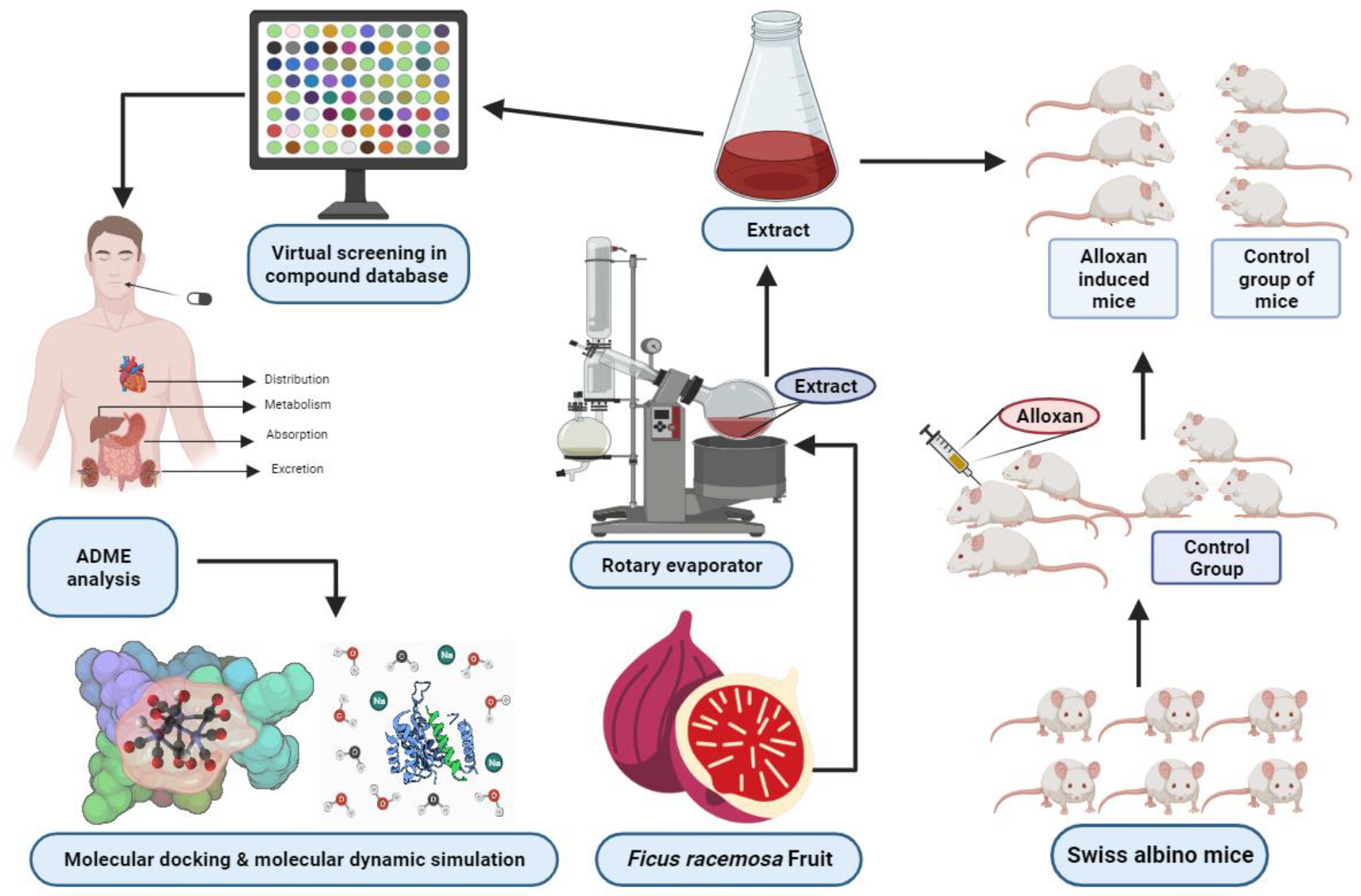
Schematic diagram of the whole study.

**Figure 2:**
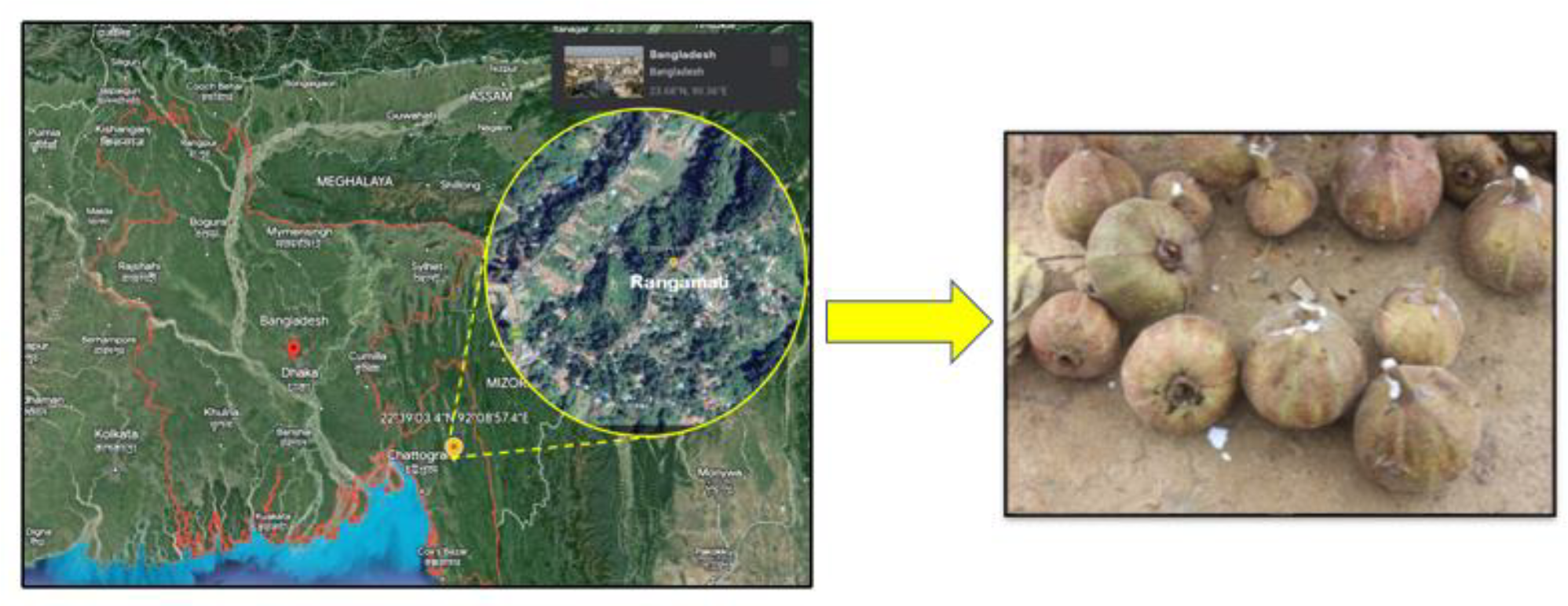
Location of the *F. racemosa s*ample collection.

### 3.1 Effect of *F. racemosa* on OGTT

Table 1 displays the effects of *F. racemosa* extract on OGTT. Before extract delivery (0 min), the BGL of all groups did not appear to differ from one another. However, all groups experienced a substantial increase (*P* < 0.01) in BGL 30 minutes after extract administration (one hour after oral glucose loading), indicating the induction of hyperglycemia. At 60 minutes, the NC group’s hyperglycemia following the glucose test had not considerably decreased, but at 120 minutes, there had been a significant change (*P* < 0.01). In contrast, compared with the glucose level at 30 minutes, all dose groups of the plant extract decreased the BGL level after 60 minutes (*P* < 0.001) and 120 minutes (*P* < 0.0001). At 60 and 120 minutes, the GC group also significantly reduced hyperglycemia (*P* < 0.001). Overall, all plant extract and GC dose groups have shown the capacity to lower BGL at 120 minutes (*P* < 0.001) (**Table 1**).

**Table 1:** The effects of *F. racemosa* fruit methanolic extract on OGTT in mice.

| Group | 0 min | 30 min | 60 min | 120 min |
| --- | --- | --- | --- | --- |
| NC | 104 ±4.5 | 176 ±8.3<sup>x**</sup> | 126 ±3.7y* | 112 ±1.2 |
| DC | 208 ±14.6 | 276 ±17.9<sup>x*</sup> | 266 ±15.1 | 262 ±14.2 |
| GC | 89 ±4.2 | 169 ±3.9<sup>x**</sup> | 99 ±2.2<sup>y*z*</sup> | 79 ±2.7<sup>x*y****</sup> |
| FR300 | 87 ±3.8 | 149 ±4.9<sup>x**</sup> | 94 ±4.4<sup>y*z*</sup> | 81 ±3.2<sup>x*y***</sup> |
| FR500 | 92 ±4.3 | 162 ±6.2<sup>x**</sup> | 88 ±3.3<sup>y*z*</sup> | 83 ±4.3<sup>x*y***</sup> |

The units of measurement are mean ± standard error. NC is the normal control group, which is given 10 milliliters per kilogram of normal saline; DC is the diabetic control that received the vehicle; GC is the positive control that receives glibenclamide drug at 5 mg/kg; FR300 and FR500 are the treatment groups that receive *F. racemosa* extract at 300 and 500 mg/kg; DS is the treatment group that receives glibenclamide at 5 mg/kg; x is compared to BGL at 0 min; y is compared to BGL after 30 min; z is compared to NC; v is compared with GC; w is compared with DC; \* $P < 0.05$ , \*\* $P < 0.01$ , \*\*\* $P < 0.001$ , \*\*\*\* $P < 0.0001$ . Blood glucose levels, or BGL. The time after extract administration is referred to as time.

### 3.2 Antihyperglycemic Activity of *F. racemosa* on Alloxan-Induced Diabetic Mice

The study assessed *F. racemosa* extract’s ability to lower blood sugar levels in diabetic mice compared to glibenclamide, a common diabetes medication. There was no statistically significant difference observed in the baseline blood glucose levels among any of the groups of mice with diabetes (**Table 2**). The results revealed that on the 7^th^ day of treatment, both the FR300 (P < 0.01) and the FR500 (P < 0.01) groups exhibited a significant decrease in blood sugar levels compared to the diabetic control group. This finding suggests that the administration of *F. racemosa* extract resulted in a significant reduction in blood glucose levels within these experimental cohorts. Similarly, in comparison to the diabetic control group, the BGL of the glibenclamide-treated group was also observed to be significantly lower on the 7^th^ (P < 0.01) and 14^th^ (P < 0.01) days of medication. Moreover, when comparing the plant extract-treated groups to the glibenclamide-treated group, no consistent statistically significant difference in blood sugar levels was observed. Furthermore, there was no significant difference in blood sugar levels between the two different doses of the plant extract (FR300 and FR500). However, the diabetic and normal control groups did not show a notable change in blood sugar levels when compared to their respective baseline levels on the seventh day of the study (**Table 2**).

**Table 2:** The effect of methanolic extract in mice.

| Group | Baseline | 7<sup>th</sup> Day |
| --- | --- | --- |
| NC | 87 ±4.5 | 85 ±3.2.3 |
| DC | 284 ±14.6 | 286 ±17.9 |
| GC | 289 ±14.2 | 79 ±3.9<sup>x**</sup> |
| FR300 | 287 ±13.8 | 111 ±4.9<sup>x**</sup> |
| FR500 | 292 ±17.3 | 86 ±6.2<sup>x**</sup> |

The units of measurement are mean ± standard error. NC is the normal control group, which is given 10 milliliters per kilogram of normal saline. DC is the diabetic control group that receives the vehicle. GC is the positive control group that receives glibenclamide medication at a dose of 5 mg/kg; FR300 and FR500 are the treatment groups that receive *F. racemosa* extract at a dose of 300 and 500 mg/kg; DS is the treatment group that receives glibenclamide at a dose of 5 mg/kg; a is compared to DC, b is compared to GC, and c is compared to baseline BGL. \**P* < 0.05, \*\**P* < 0.01, \*\*\**P* < 0.001, \*\*\**p* < 0.0001; BGL, blood glucose level.

### 3.3 Retrieval of active compounds of *F. racemosa* fruit

*F. racemosa* fruit, commonly known as fig fruit, contains several groups of compounds like phenolics, sterols, carotenoids, and organic acids. The therapeutic possibilities of *F. racemosa* is considered in various activities including hepatoprotective, anti-microbial, anti-diabetic, anti-inflammatory.

The bioactive compounds list of fig fruit was retrieved from different literatures. The phytochemical screening disclosed 13 fig fruit active compounds that have therapeutic possibilities (**Supplementary Table 1**).

### 3.4 ADME and Toxicity analysis

All available literature and databases were searched to identify active compounds from the fruit part of *F. racemosa*, resulting in the selection of 13 compounds (**Supplementary Table 2 and Supplementary Fig 1**). Subsequently, ADMET profile analysis was conducted for each of the 13 compounds to evaluate their pharmacokinetic and pharmacodynamic properties. The tool known as ADMETlab 2.0 was utilized for the purpose of conducting an analysis of adsorption, metabolism, and excretion. In addition to that, the pkCSM and Pro Tox II tools were utilized in order to analyze the distribution and toxicity, respectively. These analyses involve several properties, each of which has precise values that indicate the quality of the final result. After taking those values into consideration, additional research was carried out to screen all the compounds and choose the molecule that would serve as the benchmark for the subsequent steps. Our research concluded that only four of these compounds should be used for molecular docking: lupeol acetate, ferulic acid, friedelin, and gluanol. This decision was reached after each of these results was examined in detail, ensuring that the chosen molecules would provide the most accurate and reliable results.

The pharmacophore and QSAR properties of the compounds were analyzed for their drug likeness, drug score, toxicity, structural polarity, and oral bioavailability by using various software mentioned Section/above and the results are listed in **Tables 3 and 4**.

**Table 3:** ADMET properties of the selected four compounds.

| ADMET Properties | Properties | Ferulic acid | Friedelin | Gluanol | Lupeol acetate |
| --- | --- | --- | --- | --- | --- |
| Adsorption | Human intestinal absorption | 0.03 | 0.157 | 0.01 | 0.009 |
|  | Caco-2 permeability | -4.902 | -5.159 | -4.783 | -4.949 |
|  | MDCK | 1.46E-05 | 5.88E-06 | 8.07E-06 | 8.57E-06 |
|  | F20% | 0.047 | 0.712 | 0.006 | 0.037 |
| Distribution | Blood brain Permeability | -0.239 | 0.72 | 0.711 | 0.644 |
|  | Fraction unbound in plasma | 0.343 | 0 | 0 | 0 |
|  | Volume of distribution (Vd) (L/kg) | -1.367 | -0.272 | 0.58 | -0.12 |
|  | CNS permeability | -2.612 | -1.555 | -1.91 | -1.754 |
| Metabolism | CYP1A2-inh | 0.059 | 0.02 | 0.026 | 0.021 |
|  | CYP1A2-sub | 0.478 | 0.63 | 0.521 | 0.406 |
|  | CYP2C9-inh | 0.142 | 0.061 | 0.117 | 0.07 |
|  | CYP2C9-sub | 0.367 | 0.39 | 0.736 | 0.392 |
| Excretion | CL | 7.48 | 19.969 | 14.915 | 7.779 |
|  | T 1/2 | 0.926 | 0.033 | 0.007 | 0.009 |
| Toxicity | Hepatotoxicity | Inactive | Inactive | Inactive | Inactive |
|  | Carcinogenicity | Inactive | Inactive | Inactive | Inactive |
|  | Mutagenicity | Inactive | Inactive | Inactive | Inactive |
|  | Cytotoxicity | Inactive | Inactive | Inactive | Inactive |

**Table 4:**
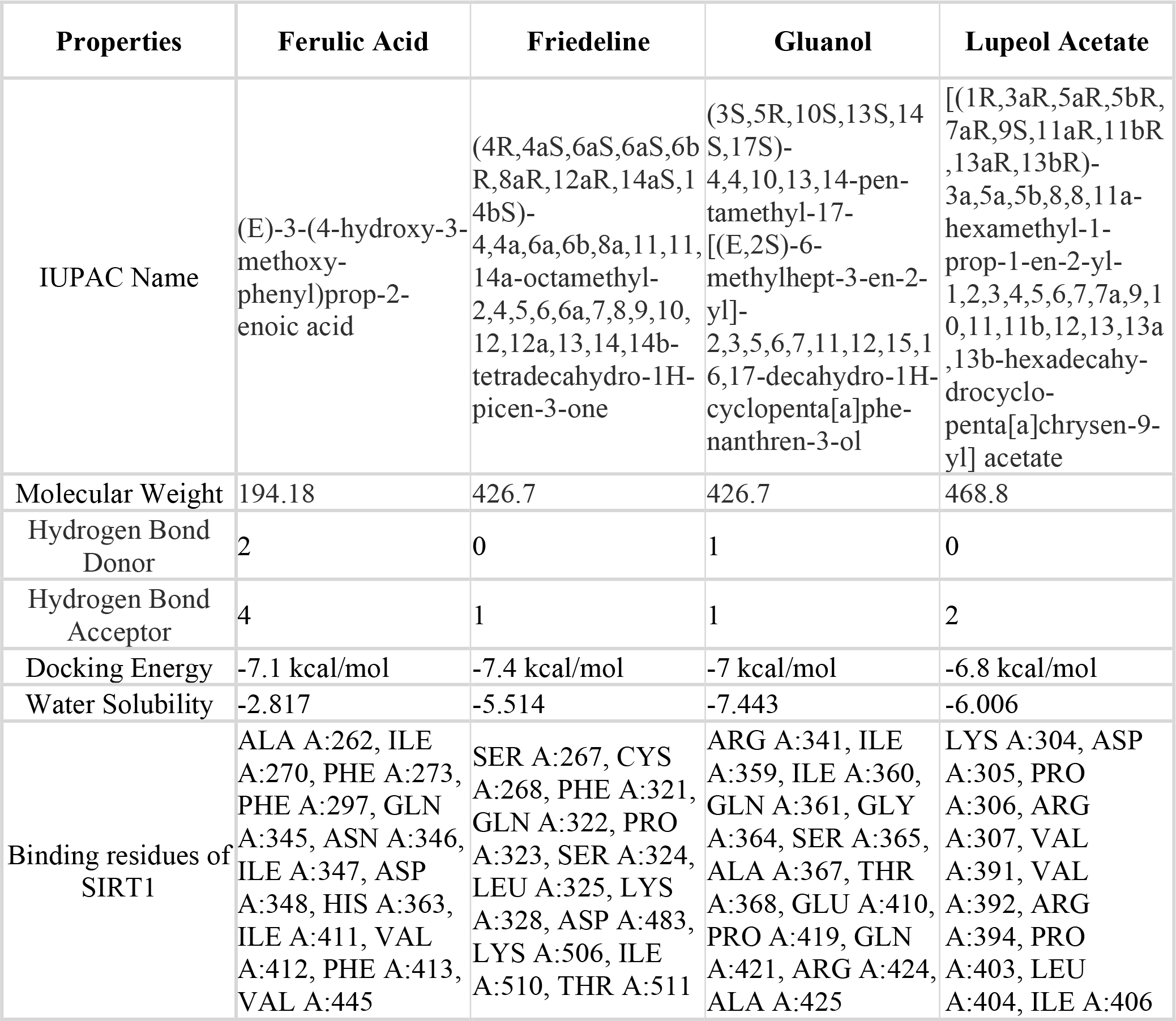
QSAR Properties of the se selected four compounds.

| Properties | Ferulic Acid | Friedelene | Gluanol | Lupeol Acetate |
| --- | --- | --- | --- | --- |
| IUPAC Name | (E)-3-(4-hydroxy-3-methoxy-phenyl)prop-2-enoic acid | (4R,4aS,6aS,6aS,6bR,8aR,12aR,14aS,14bS)-4,4a,6a,6b,8a,11,11,14a-octamethyl-2,4,5,6,6a,7,8,9,10,12,12a,13,14,14b-tetradecahydro-1H-picen-3-one | (3S,5R,10S,13S,14S,17S)-4,4,10,13,14-pentamethyl-17-[(E,2S)-6-methylhept-3-en-2-yl]-2,3,5,6,7,11,12,15,16,17-decahydro-1H-cyclopenta[a]phenanthren-3-ol | [(1R,3aR,5aR,5bR,7aR,9S,11aR,11bR,13aR,13bR)-3a,5a,5b,8,8,11a-hexamethyl-1-prop-1-en-2-yl-1,2,3,4,5,6,7,7a,9,10,11,11b,12,13,13a,13b-hexadecahydrocyclopenta[a]chrysen-9-yl] acetate |
| Molecular Weight | 194.18 | 426.7 | 426.7 | 468.8 |
| Hydrogen Bond Donor | 2 | 0 | 1 | 0 |
| Hydrogen Bond Acceptor | 4 | 1 | 1 | 2 |
| Docking Energy | -7.1 kcal/mol | -7.4 kcal/mol | -7 kcal/mol | -6.8 kcal/mol |
| Water Solubility | -2.817 | -5.514 | -7.443 | -6.006 |
| Binding residues of SIRT1 | ALA A:262, ILE A:270, PHE A:273, PHE A:297, GLN A:345, ASN A:346, ILE A:347, ASP A:348, HIS A:363, ILE A:411, VAL A:412, PHE A:413, VAL A:445 | SER A:267, CYS A:268, PHE A:321, GLN A:322, PRO A:323, SER A:324, LEU A:325, LYS A:328, ASP A:483, LYS A:506, ILE A:510, THR A:511 | ARG A:341, ILE A:359, ILE A:360, GLN A:361, GLY A:364, SER A:365, ALA A:367, THR A:368, GLU A:410, PRO A:419, GLN A:421, ARG A:424, ALA A:425 | LYS A:304, ASP A:305, PRO A:306, ARG A:307, VAL A:391, VAL A:392, ARG A:394, PRO A:403, LEU A:404, ILE A:406 |

Among the ADMET characteristics, human intestinal absorption of lupeol acetate (0.009 out of 1.0) is the lowest, whereas Caco-2 permeability of Friedelin is the lowest at -5.159. All the other compounds are of high quality for human intestine absorption (0.01; 0.03; 0.157); and Caco-2 permeability, with all values exceeding -5.15. In terms of MDCK permeability, the optimal result is greater than 2 x 10-6, indicating that ferulic acid has an exceptional quality of 1.46 x 10-05. However, the results for the remaining three compounds are still impressive. Friedelin has the lowest oral bioavailability score of the four substances, 0.712, as a score between 0.7 and 1.0 indicates poor quality. The results for the remaining three compounds range from good to acceptable (**Table 3**).

Friedelin had the best result for the blood-brain barrier, with a revealed score of 0.72, while ferulic acid had the worst, with a revealed score of -0.239. Results for fraction unbound in plasma satisfies the standard level for all four compounds. It should be noted that three of the four compounds performed poorly in terms of volume of distribution (Vd), with only gluanol scoring at the standard level with a score of 0.58. CNS permeability, the good quality result stands with logPS > -2 as it is considered to penetrate the central nervous system. All compounds except ferulic acid showed good quality results (-1.91 --1.555).

The output value for metabolism is the probability of being a substrate or inhibitor, between 0 and 1. All CYP1A2 and CYP2C9 inhibitors exhibited positive findings, as their ranges are significantly lower than 1. In the case of these isozyme substrates, however, all results are quite modest, with the exception of Gluanol, whose CYP2C9-sub score is 0.736, which is very close to 1.

The standard value for a drug’s clearance is at least 5. All compounds produced an outstanding outcome for CL. In contrast, the half-lives of these chemicals are excellent, with the exception of ferulic acid, which scored 0.926. The chart of toxicity data revealed that none of the compounds exhibited any action when tested for hepatotoxicity, carcinogenicity, mutagenicity, or cytotoxicity (**Table 3**).

### 3.5 Molecular docking and Interpretation of protein-ligands interactions

The SIRT1 protein was modeled and the active site of the protein was determined before the docking experiments (**Supplementary Fig 2 and 3**). Four compounds (Friedelin, Lupeol Acetate, Gluanol, and Ferulic Acid) were docked with SIRT1 using AutoDock Vina to determine their binding affinity (**Fig 3**). AutoDock Vina generated nine potential binding locations, and the optimal one was picked for each molecule based on the lowest docking energy and binding amino acid residues (**Fig 4 and Supplementary Fig 4**). The binding affinity of friedelin, ferulic acid, gluanol, and lupeol acetate with the protein SIRT1 was -7.4 kcal/mol, -7.1 kcal/mol, -7 kcal/mol, and -6.8 kcal/mol, respectively. Friedelin formed one conventional hydrogen bond with LEU A:325, ten van der Waals interactions with SER A:267, CYS A:268, GLN A:322, PRO A:323, SER A:324, LYS A:328, ASP A:483, LYS A:506, ILE A:510, THR A:511, and one Pi-alkyl bonds with PHE A:321, with the SIRT1 protein after performing molecular docking (**Fig 4A**). Lupeol acetate exhibited one conventional hydrogen bonds with ARG A:307, four van der Waals interactions with LYS A:304, ASP A:305, LEU A:404, ILE A:406, and five alkyl bonds with PRO A:306, VAL A:391, VAL A:392, ARG A:394, PRO A:403 (**Fig 4B**). Van der Waals bonds were predominantly formed with six amino acid residues, for Gluanol, including ARG A:341, GLN A:361, GLY A:364, SER A:365, GLN A:421, ARG A:424, two conventional hydrogen bonds with THR A:368, GLU A:410, and five alkyl bonds with ILE A:359, ILE A:360, ALA A:367, PRO A:419, ALA A:425 (**Fig 4C**). Ferulic acid, the fourth compound, had five conventional hydrogen bonds at ASN A:346, ILE A:347, ASP A:348, HIS A:363, VAL A:412, six van der Waals interactions with ALA A:262, ILE A:270, GLN A:345, ILE A:411, PHE A:413, VAL A:445, one alkyl bonds with ILE A:347, and two Pi-Pi T-shaped at PHE A:273, PHE A:297 (**Fig 4D**).

**Figure 3:**
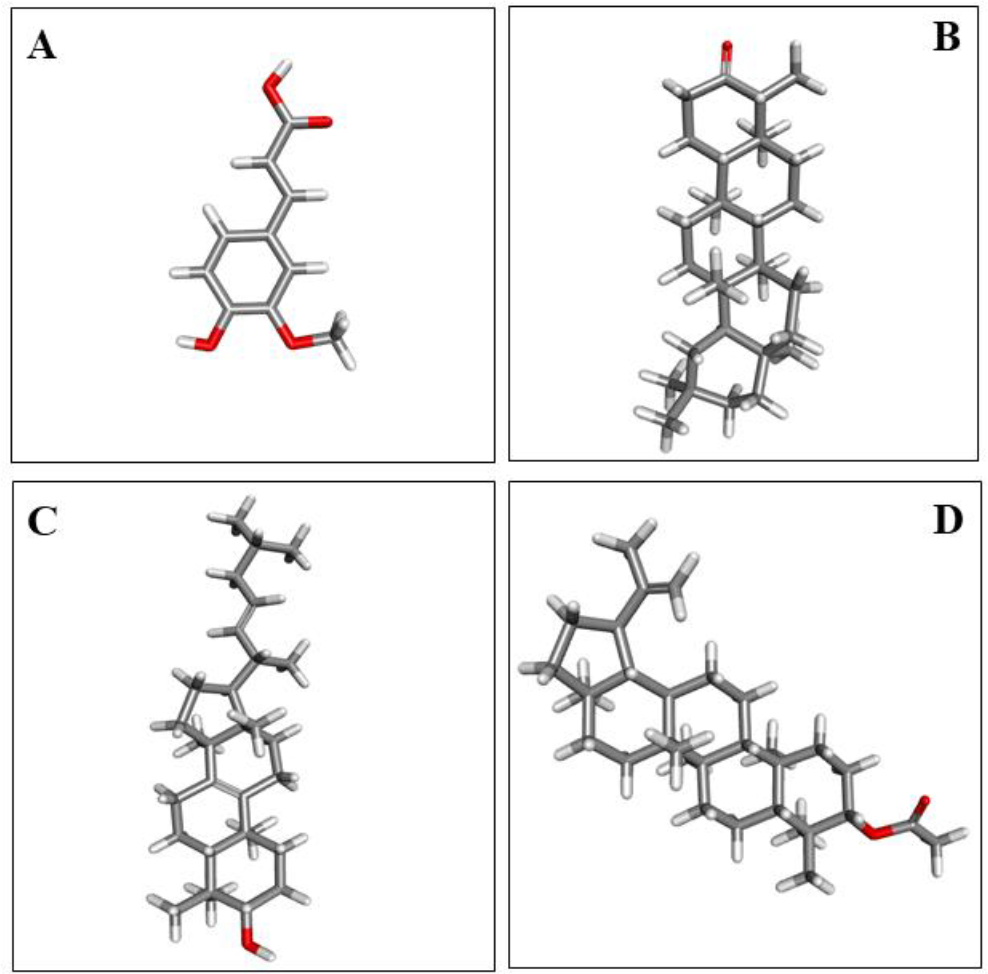
Two-dimensional structures of the four candidate drugs from *Ficus racemosa*. (A) Ferulic acid, (B) Friedelin, (C) Gluanol, (D) Lupeol acetate.

### 3.6 Molecular Dynamics Simulation

Analysis of the docking complex of our four protein ligands indicate acceptable deviations of 0.3 nm, with one notable exception of SIRT1-Friedeline complex. The SIRT1-Friedeline complex demonstrated a maximum variation of 0.3 nm between 25 and 35 ns during a simulation spanning 100 ns. The plot indicated that after 50 ns, the entire SIRT1-drug complex obtained stability and, in the last 30 ns, became concurrent with the SIRT1 complex’s RMSD plot (**Fig 5A**). The RMSF value for all SIRT1 complex residues is minimal and oscillates between 0 and 0.5 nm, with the exception of the minimum of 0.1 nm with a maximum of 2.25 nm in residues from 500 until the end residue of the protein-ligand complex. The plot’s value explains that the 250–500 residue region with an RMSF value of less than 0.5 nm is not very flexible nor dynamic (**Fig 5B**). The complexes demonstrate that the radius of gyration (Rg) of all four (4) demonstrated fluctuations in the 100 ns time frame within the permissible range of 0.2 nm with little variations; however, the SIRT1-Gluanol complex best preserved the sturdy and linear without much fluctuation or variation. The SIRT1-Friedeline complex presented high fluctuations and variations during the period (**Fig 5C**). The protein-ligand complexes’ SASA values exhibited a consistent pattern over the course of 100 ns, however all of them showed fluctuations in the first 20 seconds, except for the SIRT1-Friedeline complex, which gradually increased. The SIRT1-Ferulic acid combination displayed the highest SASA value over the course of 100 ns, with the other three complexes showing SASA values that gradually decreased after 30 ns and the SIRT1-Friedeline complex showing minor variations. The SIRT1-Gluanol and SIRT1-Lupeol acetate complexes were the only ones to consistently maintain the SASA value with few changes (**Fig 5D**).

**Figure 5:**
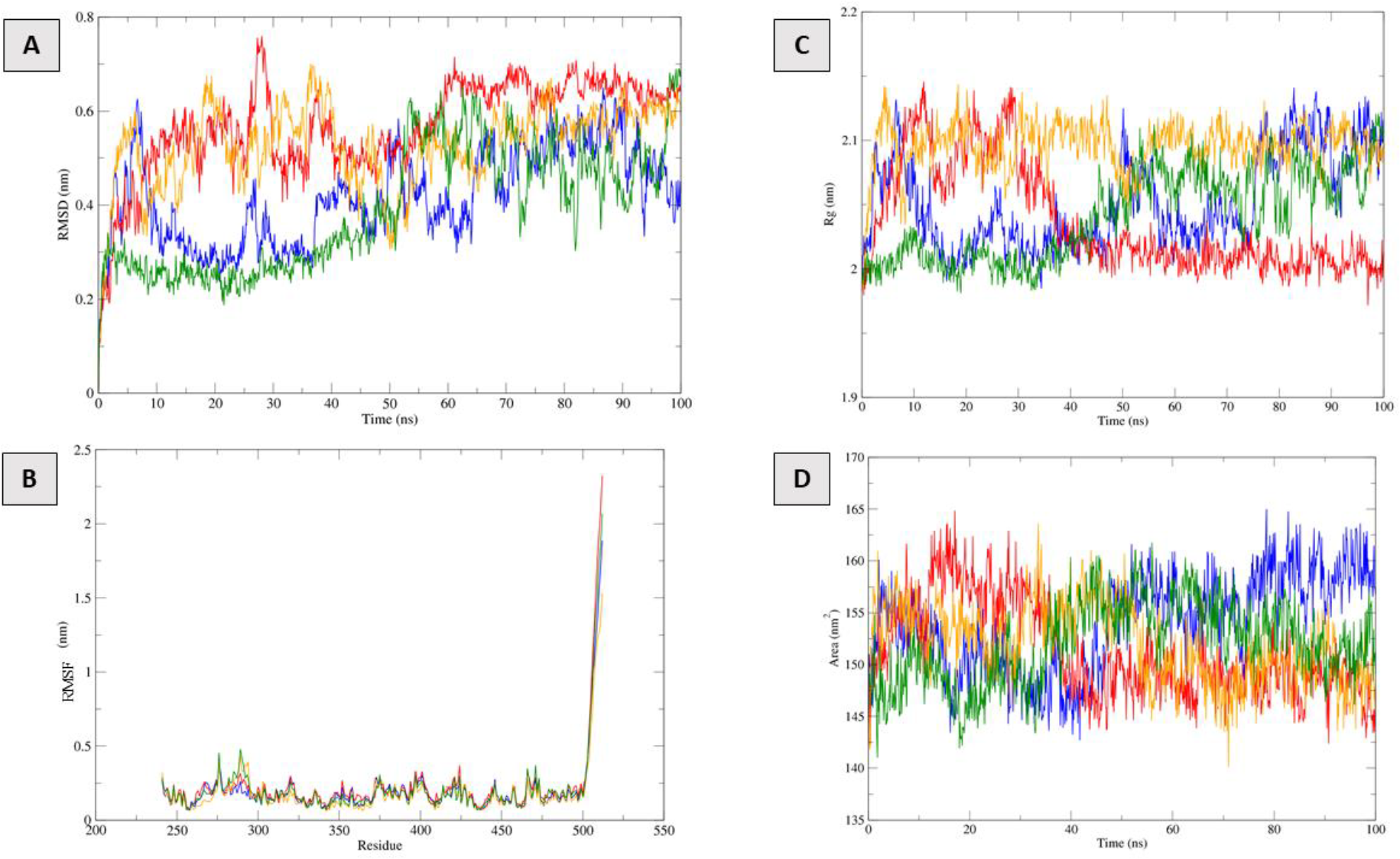
Molecular dynamics simulation of the SIRT1 and ligand complex, A. RMSD; B. RMSF C. Rg; D. SASA, viz SIRT1-Ferulic acid complex (Blue); SIRT1-Friedeline complex (Red); SIRT1-Gluanol complex (Orange); SIRT1-Lupeol acetate complex (Green).

## 4. Discussion

The present study aims to investigate the antidiabetic effect of the *F. racemosa* plant extract on experimental diabetic mice and exploring the antidiabetic potentials of the compounds of *F. racemosa*. In addition, the study examined the potential interaction between these compounds and the protein associated with diabetes using molecular docking and molecular dynamics (MD) simulations, demonstrating their potential as a natural alternative to drugs for treating diabetes.

Initially, a dosage of 150 mg/kg of alloxan was administered to both male and female mice, leading to the suppression of glucokinase activity and subsequent development of chronic hyperglycemia in the mice (41,42). Afterwards, the safe dose for the extract was experimented and the findings suggested that any dose under 3000 mg/kg is safe to be used in mice. Similar dose has been also documented in the following study (43). The results of the oral glucose tolerance test (OGTT) indicated that all doses of the plant extract exhibited the capacity to effectively utilize glucose within the body. The extract, in both doses, has the potential to inhibit the absorption of glucose in the intestines and liver, thereby leading to a reduction in blood glucose levels (44,45). Nevertheless, the diabetic mice exhibited impaired glucose utilization, potentially attributable to beta cell dysfunction and reduced glucose uptake capacity (**Table 1**). Following the administration of *F. racemosa* plant extract at doses of 300 mg/kg and 500 mg/kg in female and male mice, a decrease in blood sugar levels was observed after 7 days. The stimulation of insulin release by plant parts has been documented as a factor in the induction of antihyperglycemic activity in mice (46,47). The administration of the plant extract may induce a reduction in gluconeogenesis and an increase in liver glycogen synthesis (48). The administration of plant extract may lead to a decrease in blood glucose levels due to the elevation of insulin and glycogen levels. The increase in glycated hemoglobin levels can be attributed to the higher concentration of glucose in the bloodstream. This indicates that elevated blood glucose levels are responsible for the rise in glycated hemoglobin (49). However, it has been observed that the impact of plant extract doses at 300 mg/kg and 500 mg/kg can lead to a decrease in both blood glucose levels and glycated hemoglobin. The drug glibenclamide is extensively utilized for the treatment and control of type 2 diabetes mellitus. This drug induces hypoglycemia by enhancing insulin secretion through the stimulation of beta cells (50). The analysis confirmed that the given doses 300 mg/kg and 500 mg/kg have the same result in terms of lowering the blood glucose in glibenclamide (**Table 2**).

After the confirmation of antidiabetic effect of the plant extract, virtual screening identified 13 compounds of *F. racemosa* from the PubChem database for further experiments. The compounds selected based on their reported therapeutic potential, were subjected to a series of computational analyses to assess their pharmacokinetic and toxicity properties, as well as their ability to bind effectively with the target protein associated with diabetes (**Supplementary Table 1 and 2**).

The ADMET and toxicity analysis have provided valuable information about the pharmacokinetic and safety profiles of the selected *F. racemosa* compounds (**Table 3 and Supplementary Table 2**). The majority of the compounds exhibited high human intestinal absorption (0.01-0.157) and good Caco-2 permeability (above -5.15). The blood-brain barrier exhibits a high degree of selectivity, allowing only a few compounds, such as ferulic acid, to cross its boundaries. As a result, the visible distribution of many compounds throughout the brain may be limited. The reason behind this variation in ADMET parameters could be attributed to the differences in molecular size, polarity, and charge distribution of the compounds. In terms of metabolic stability, all compounds showed low probability of being a substrate or inhibitor of CYP1A2 and CYP2C9 enzymes, except for gluanol, which showed a high probability of being a CYP2C9 substrate. Because, the variation in metabolic stability could be attributed to the presence of functional groups and the site of metabolism in the compounds. All the compounds are determined to be ideal to be used as an oral medication following the analysis of metabolic profiling. Regarding toxicity parameters, all compounds showed no hepatotoxicity, carcinogenicity, mutagenicity, or cytotoxicity, which is a favorable result.

Overall, the ADMET and toxicity analysis results suggest that the selected *F. racemosa* compounds have favorable pharmacokinetic and safety profiles, with some compounds showing superior characteristics than others. These findings can be useful in selecting lead compounds for further drug development studies (**Table 3**).

The identification of the active site and of SIRT1 protein provides valuable insights into the protein’s function and can be utilized in the development of new therapeutics targeting SIRT1. We investigated the binding affinity of four compounds (Friedelin, Lupeol Acetate, Gluanol, and Ferulic Acid) with the protein SIRT1 using PyRx tool AutoDock Vina. We identified the optimal binding location for each molecule based on the lowest docking energy and binding amino acid residues. The binding affinity of each compound with SIRT1 was determined, and we found that Friedelin, Lupeol Acetate, Gluanol, and Ferulic Acid had binding affinities of -7.4 kcal/mol, -6.8 kcal/mol, -7 kcal/mol, and -7.1 kcal/mol, respectively (**Table 4**). We also identified the specific amino acid residues that formed conventional hydrogen bonds, van der Waals interactions, and alkyl bonds with each compound. These findings suggest that these compounds may have potential therapeutic uses in the treatment of diseases associated with SIRT1 (**Fig 4**). These interactions are important for stabilizing the protein-ligand complex and determining the binding affinity.

In addition, the molecular dynamics simulations have demonstrated temporal changes in many parameters including Rg, RMSD, RMSF, and SASA during a 100-nanosecond time period. Previous research has conducted numerous molecular dynamics simulation experiments to examine the temporal stability of molecules (51,52). Nevertheless, the aforementioned results exhibited a consistent pattern throughout time, indicating a strong affinity between the chemical and the receptor (**Fig 5**). In conclusion, our study proposes four prospective antidiabetic substances for the treatment of diabetes and examines their potential impact on reducing hyperglycemia in individuals with diabetes (**Fig 6**).

**Figure 6:**
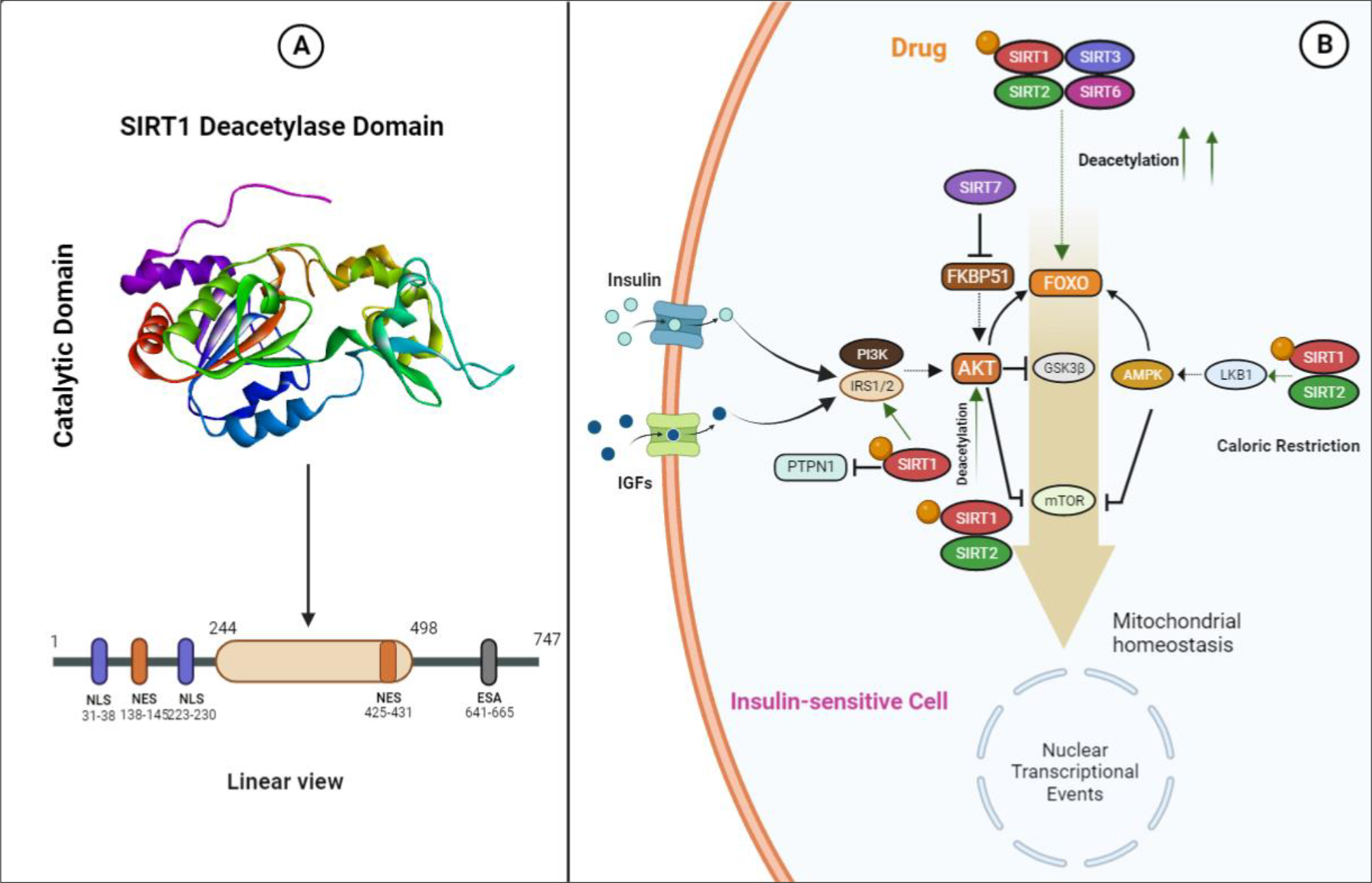
The possible antidiabetic effect of F. racemose plant extract. A) Effect of SIRT1, B) Putative insulin regulation by SIRT1-compounds interactions.

### Limitations of the study

Nonetheless, it is important to acknowledge several limitations in our study on the potential therapeutic effects of *F. racemosa* compounds on diabetes. The study focused on four compounds and did not investigate other potentially effective compounds present in the plant extract. Additionally, the study used computational models and predictions to evaluate the ADME and toxicity of the compounds, and the results may not reflect their actual pharmacokinetic and safety profiles in humans. Furthermore, the study did not investigate the long-term effects of the compounds on diabetes treatment, and further research is needed to determine their safety and efficacy. Despite these limitations, the study’s findings are promising and suggest that *F. racemosa* compounds may have potential as a treatment for diabetes, although more research is needed to fully understand their effects.

## 5. Conclusion

The present study has evaluated the antidiabetic potentials of *F. racemosa* fruit in mice model. Further, the compounds of *F. racemosa* were virtually screened and prioritized according to their ADMET profile. Based on the data, four substances seem like good candidates for possible medicinal applications. Moreover, the interactions of these drug candidates with SIRT1 receptor seemed to be stable as predicted by RMSD, RMSF, Rg, SASA. Finally, the findings of this study may explore the new promising avenue in drug discovery field.

## 8. Author Contribution

**Mohammad Uzzal Hossain, A.B.Z Naimur Rahman, Md. Shahadat Hossain, Shajib Dey**,, **Zeshan Mahmud Chowdhury**: Conceptualization; data retrieval; sequence analysis; laboratory experiments; writing – original draft; **Arittra Bhattacharjee**: writing – review and editing; Ishtiaque Ahammad: manuscript preparations; **Mohammad Kamrul Hasan**: data curation; methodology; Md. Billal Hosen: data curation; methodology; Istiak Ahmed: data curation; methodology; **Keshob Chandra Das**: supervision; validation; **Chaman Ara Keya**: supervision; validation; **Md. Salimullah**: project administration; validation; supervision.

## References

1. American Diabetes Association. 2. Classification and Diagnosis of Diabetes: Standards of Medical Care in Diabetes—2021. Diabetes Care.2021 Jan 1;44(Supplement_1):S15–33.

2. ElSayed NA, Aleppo G, Aroda VR, Bannuru RR, Brown FM, Bruemmer D, et al. 8. Obesity and Weight Management for the Prevention and Treatment of Type 2 Diabetes: Standards of Care in Diabetes—2023. Diabetes Care.2023 Jan 1;46(Supplement_1):S128–39.

3. Kahn SE, Cooper ME, Del Prato S. Pathophysiology and treatment of type 2 diabetes: perspectives on the past, present, and future. The Lancet. 2014 Mar;383(9922):1068–83.

4. National Institute of Diabetes and Digestive and Kidney Diseases. “Diabetes Diet, Eating, & Physical Activity” [Internet]. 2023 [cited 2023 Oct 12]. Available from: https://www.niddk.nih.gov/health-information/diabetes/overview/diet-eating-physical-activity

5. van Baar MJB, van Ruiten CC, Muskiet MHA, van Bloemendaal L, IJzerman RG, van Raalte DH. SGLT2 Inhibitors in Combination Therapy: From Mechanisms to Clinical Considerations in Type 2 Diabetes Management. Diabetes Care.2018 Aug 1;41(8):1543–56.

6. Ong KL, Stafford LK, McLaughlin SA, Boyko EJ, Vollset SE, Smith AE, et al. Global, regional, and national burden of diabetes from 1990 to 2021, with projections of prevalence to 2050: a systematic analysis for the Global Burden of Disease Study 2021. The Lancet. 2023 Jul;402(10397):203–34.

7. International Diabetes Federation. International Diabetes Federation. 2023 [cited 2023 Oct 17]. Diabetes Facts and Figures. Available from: https://idf.org/about-diabetes/diabetes-factsfigures/

8. Statista. Statista. 2021 [cited 2023 Oct 17]. Number of diabetics worldwide by region 2021. Available from: https://www.statista.com/statistics/241802/number-of-diabetics-worldwide-by-region/

9. Galicia-Garcia U, Benito-Vicente A, Jebari S, Larrea-Sebal A, Siddiqi H, Uribe KB, et al. Pathophysiology of Type 2 Diabetes Mellitus. Int J Mol Sci.2020 Aug 30;21(17).

10. Williams R, Karuranga S, Malanda B, Saeedi P, Basit A, Besançon S, et al. Global and regional estimates and projections of diabetes-related health expenditure: Results from the International Diabetes Federation Diabetes Atlas, 9th edition. Diabetes Res Clin Pract. 2020 Apr;162:108072.

11. World Health Organization. World Health Organization. 2023 [cited 2023 Oct 17]. Diabetes. Available from: https://www.who.int/news-room/fact-sheets/detail/diabetes

12. Kahn SE, Cooper ME, Del Prato S. Pathophysiology and treatment of type 2 diabetes: perspectives on the past, present, and future. The Lancet. 2014 Mar;383(9922):1068–83.

13. Standards of Medical Care in Diabetes—2021 Abridged for Primary Care Providers. Clinical Diabetes.2021 Jan 1;39(1):14–43.

14. Ahmed F, Urooj A. Traditional uses, medicinal properties, and phytopharmacology of Ficus racemosa : A review. Pharm Biol.2010 Jun 6;48(6):672–81.

15. Grover JK, Yadav S, Vats V. Medicinal plants of India with anti-diabetic potential. J Ethnopharmacol. 2002 Jun;81(1):81–100.

16. Walia A, Kumar N, Singh R, Kumar H, Kumar V, Kaushik R, et al. Bioactive Compounds in Ficus Fruits, Their Bioactivities, and Associated Health Benefits: A Review. J Food Qual.2022 Apr 26;2022:1–19.

17. Amin MM, Bhakta S, Das SK. Anti-diabetic potential of Ficus racemosa: current state and prospect especially in the developing countries. Journal of Bioscience and Agriculture Research. 2015;5(2):65–72.

18. Chaware GK, Kumar V, Kumar S, Kumar P. Bioactive Compounds, Pharmacological Activity and Food Application of Ficus racemosa : A Critical Review. International Journal of Fruit Science.2020 Sep 14;20(sup2):S969–86.

19. Deepa P, Sowndhararajan K, Kim S, Park SJ. A role of Ficus species in the management of diabetes mellitus: A review. J Ethnopharmacol. 2018 Apr;215:210–32.

20. Kumar GPS, Arulselvan P, Kumar DS, Subramanian SP. Anti-Diabetic Activity of Fruits of Terminalia chebula on Streptozotocin Induced Diabetic Rats. Journal of Health Science. 2006;52(3):283–91.

21. Miura T, Koike T, Ishida T. Antidiabetic Activity of Green Tea (Thea sinensis L.) in Genetically Type 2 Diabetic Mice. Journal of Health Science. 2005;51(6):708–10.

22. Nagappa AN, Thakurdesai PA, Venkat Rao N, Singh J. Antidiabetic activity of Terminalia catappa Linn fruits. J Ethnopharmacol. 2003 Sep;88(1):45–50.

23. National Research Council (U.S.). Committee for the Update of the Guide for the Care and Use of Laboratory Animals., Institute for Laboratory Animal Research (U.S.). Guide for the care and use of laboratory animals. National Academies Press; 2011. 220 p.

24. Ragavan B, Krishnakumari S. Antidiabetic effect of T. arjuna bark extract in alloxan induced diabetic rats. Indian Journal of Clinical Biochemistry. 2006 Sep;21(2):123–8.

25. Oecd. OECD GUIDELINE FOR TESTING OF CHEMICALS Acute Oral Toxicity-Up-and-Down Procedure INTRODUCTION. 2001.

26. Bürger C, Fischer DR, Cordenunzzi DA, Batschauer AP de B, Cechinel Filho V, Soares AR dos S. Acute and subacute toxicity of the hydroalcoholic extract from Wedelia paludosa (Acmela brasiliensis) (Asteraceae) in mice. J Pharm Pharm Sci.2005 Aug 19;8(2):370–3.

27. Carvalho EN de, Carvalho NAS de, Ferreira LM. Experimental model of induction of diabetes mellitus in rats. Acta Cir Bras. 2003;18(spe):60–4.

28. Kamalakkannan N, Prince PSM. Hypoglycaemic effect of water extracts of Aegle marmelos fruits in streptozotocin diabetic rats. J Ethnopharmacol. 2003 Aug;87(2–3):207–10.

29. Gidado A, Ameh DA, Atawodi SE. Effect of Nauclea latifolia leaves aqueous extracts on blood glucose levels of normal and alloxan-induced diabetic rats. Afr J Biotechnol. 2005;4(1):91–3.

30. Pari L, Venkateswaran S. Effect of an aqueous extract of Phaseolus vulgaris on the properties of tail tendon collagen of rats with streptozotocin-induced diabetes. Brazilian Journal of Medical and Biological Research. 2003 Jul;36(7):861–70.

31. Berman HM. The Protein Data Bank. Nucleic Acids Res.2000 Jan 1;28(1):235–42.

32. Kim S, Chen J, Cheng T, Gindulyte A, He J, He S, et al. PubChem in 2021: new data content and improved web interfaces. Nucleic Acids Res.2021 Jan 8;49(D1):D1388–95.

33. Xiong G, Wu Z, Yi J, Fu L, Yang Z, Hsieh C, et al. ADMETlab 2.0: an integrated online platform for accurate and comprehensive predictions of ADMET properties. Nucleic Acids Res.2021 Jul 2;49(W1):W5–14.

34. Pires DE V., Blundell TL, Ascher DB. pkCSM: Predicting Small-Molecule Pharmacokinetic and Toxicity Properties Using Graph-Based Signatures. J Med Chem.2015 May 14;58(9):4066–72.

35. Johansson MU, Zoete V, Michielin O, Guex N. Defining and searching for structural motifs using DeepView/Swiss-PdbViewer. BMC Bioinformatics.2012 Dec 23;13(1):173.

36. Hanwell MD, Curtis DE, Lonie DC, Vandermeersch T, Zurek E, Hutchison GR. Avogadro: an advanced semantic chemical editor, visualization, and analysis platform. J Cheminform.2012 Dec 13;4(1):17.

37. Dallakyan S, Olson AJ. Small-Molecule Library Screening by Docking with PyRx. In 2015. p. 243–50.

38. Schrödinger LLC, DeLano W. PyMOL [Internet]. Available from: http://www.pymol.org/pymol

39. SYSTÉMES D. 2016. “BIOVIA Discovery Studio” “Dassault Syst mes BIOVIA, Discovery Studio Modeling Environment, Release 2017” Dassault Syst mes .

40. Schuler LD, Daura X, van Gunsteren WF. An improved GROMOS96 force field for aliphatic hydrocarbons in the condensed phase. J Comput Chem. 2001 Aug;22(11):1205–18.

41. Rohilla A, Ali S. Alloxan Induced Diabetes : Mechanisms and Effects. International Journal of Research in Pharmaceutical and Biomedical Science. 2012;3(2).

42. Radenković M, Stojanović M, Prostran M. Experimental diabetes induced by alloxan and streptozotocin: The current state of the art. J Pharmacol Toxicol Methods. 2016 Mar;78:13–31.

43. Amare YE, Dires K, Asfaw T. Antidiabetic Activity of Mung Bean or Vigna radiata (L.) Wilczek Seeds in Alloxan-Induced Diabetic Mice. Evidence-Based Complementary and Alternative Medicine.2022 Oct 26;2022:1–12.

44. Hetta MH, Owis AI, Haddad PS, Eid HM. The fatty acid-rich fraction of Eruca sativa (rocket salad) leaf extract exerts antidiabetic effects in cultured skeletal muscle, adipocytes and liver cells. Pharm Biol. 2017 Dec;55(1):810–8.

45. Naowaboot J, Pannangpetch P, Kukongviriyapan V, Prawan A, Kukongviriyapan U, Itharat A. Mulberry Leaf Extract Stimulates Glucose Uptake and GLUT4 Translocation in Rat Adipocytes. Am J Chin Med (Gard City N Y).2012 Jan 30;40(01):163–75.

46. Pari L, Latha M. Effect of Cassia auriculata flowers on blood sugar levels, serum and tissue lipids in streptozotocin diabetic rats. Singapore Med J. 2002 Dec;43(12):617–21.

47. Lee MK, Kim MJ, Cho SY, Park SA, Park KK, Jung UJ, et al. Hypoglycemic effect of Duzhong (Eucommia ulmoides Oliv.) leaves in streptozotocin-induced diabetic rats. Diabetes Res Clin Pract. 2005 Jan;67(1):22–8.

48. Khan BA, Abraham A, Leelamma S. Hypoglycemic action of Murraya koenigii (curry leaf) and Brassica juncea (mustard): mechanism of action. Indian J Biochem Biophys. 1995 Apr;32(2):106–8.

49. Pari L, Saravanan R. Antidiabetic effect of diasulin, a herbal drug, on blood glucose, plasma insulin and hepatic enzymes of glucose metabolism in hyperglycaemic rats. Diabetes Obes Metab.2004 Jul 26;6(4):286–92.

50. Vangiersbergen P, Treiber A, Clozel M, Bodin F, Dingemanse J. In vivo and in vitro studies exploring the pharmacokinetic interaction between bosentan, a dual endothelin receptor antagonist, and glyburide. Clin Pharmacol Ther. 2002 Apr;71(4):253–

51. Hossain MU, Khan MA, Rakib-Uz-Zaman SM, Ali MT, Islam MS, Keya CA, Salimullah M. Treating diabetes mellitus: pharmacophore based designing of potential drugs from Gymnema sylvestre against insulin receptor protein. BioMed research international. 2016 Oct;2016.

52. Hossain MU, Bhattacharjee A, Emon MT, Chowdhury ZM, Ahammad I, Mosaib MG, Moniruzzaman M, Rahman MH, Islam MN, Ahmed I, Amin MR. Novel mutations in NSP-1 and PLPro of SARS-CoV-2 NIB-1 genome mount for effective therapeutics. Journal of Genetic Engineering and Biotechnology. 2021 Dec;19(1):1–0.

