## Supplementary material for "Mice Models to Bioinformatics Methods Led to the Discovery of Antidiabetic Compounds in the *F. racemosa* Plant Extracts": Supplemntary file

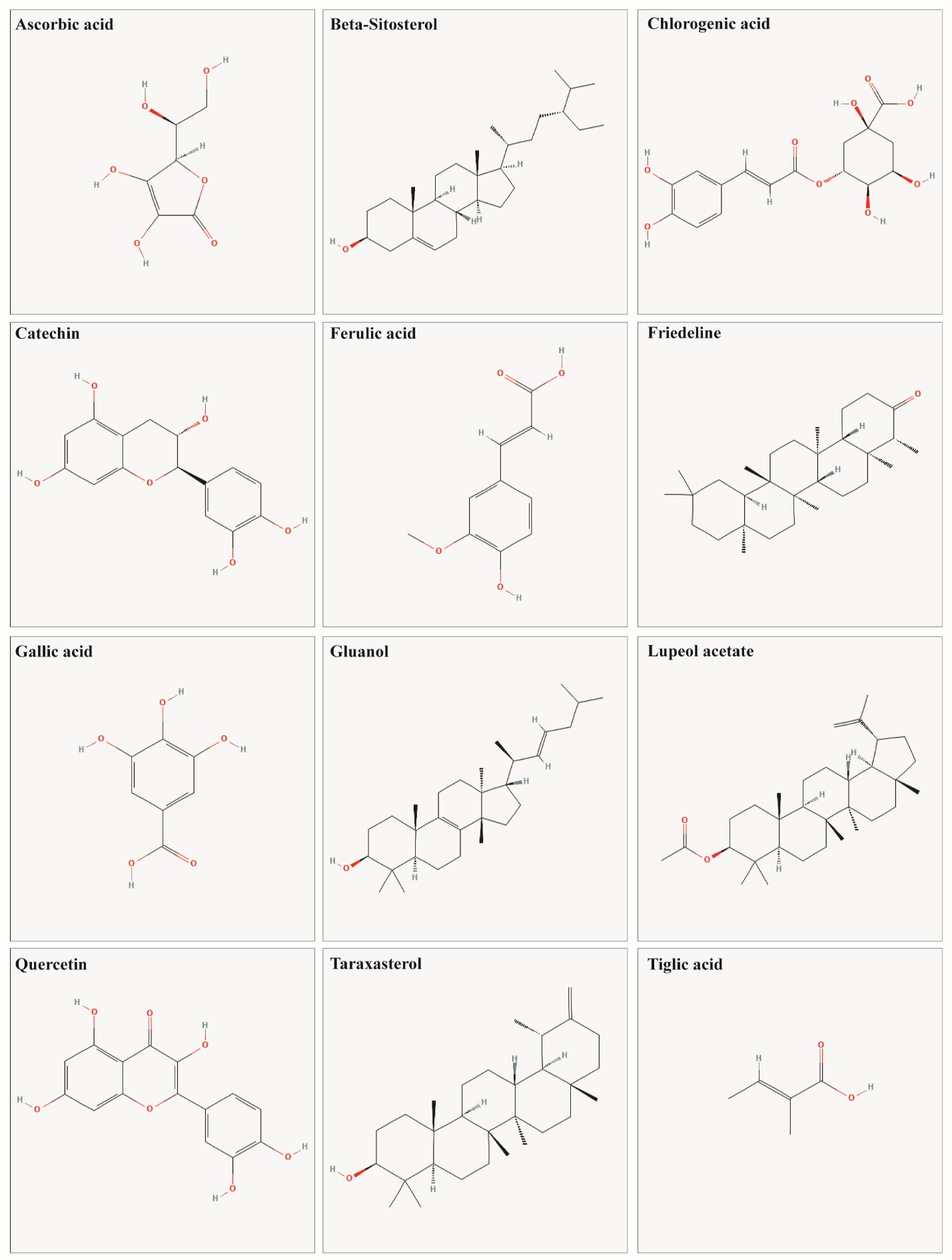

**Supplementary Figure 1:** Two-dimensional structures of all 13 compounds from *Ficus racemosa*.

**Supplementary Figure 2: Three-dimensional (3D) crystal structure of SIRT1 protein (PDB ID: 4I5I)**

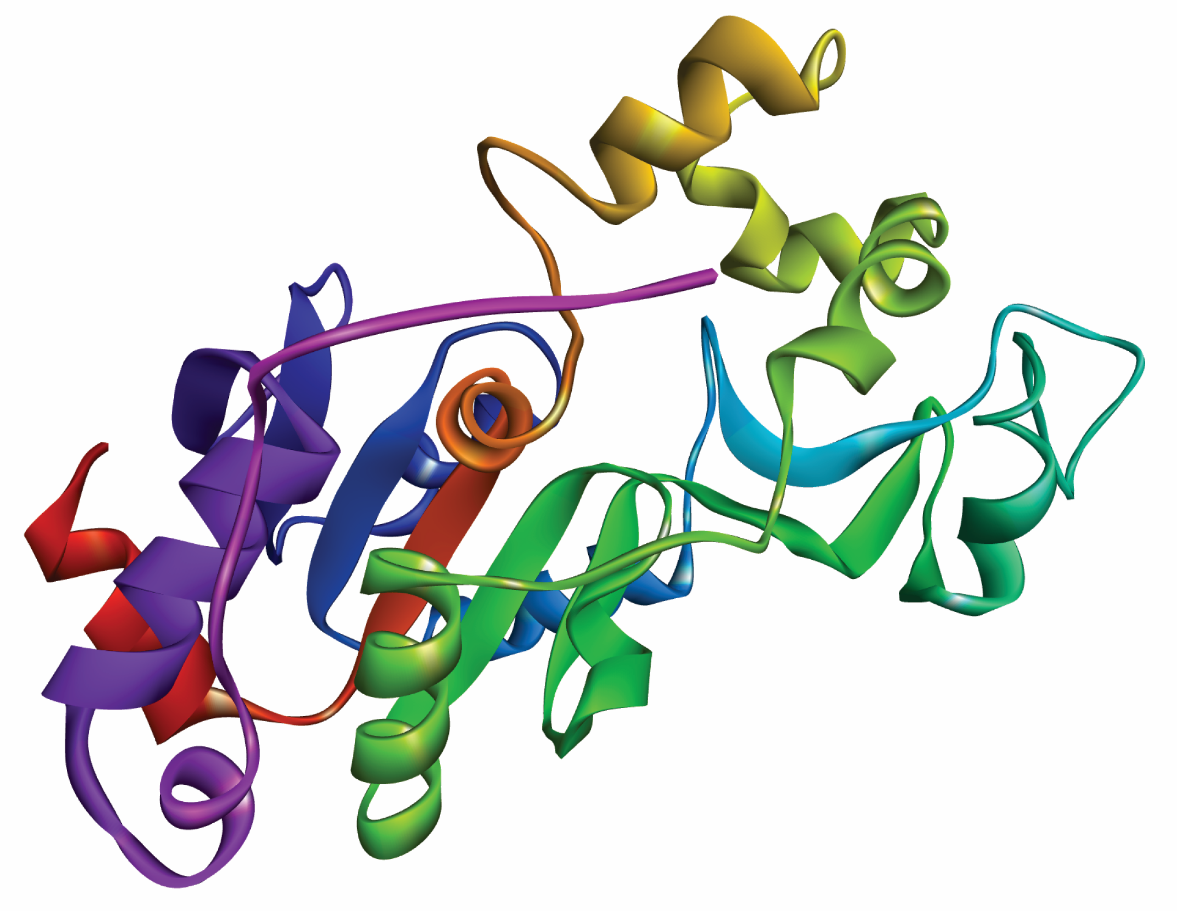

**Supplementary Figure 2:** Three-dimensional (3D) crystal structure of SIRT1 protein (PDB ID: 4I5I)

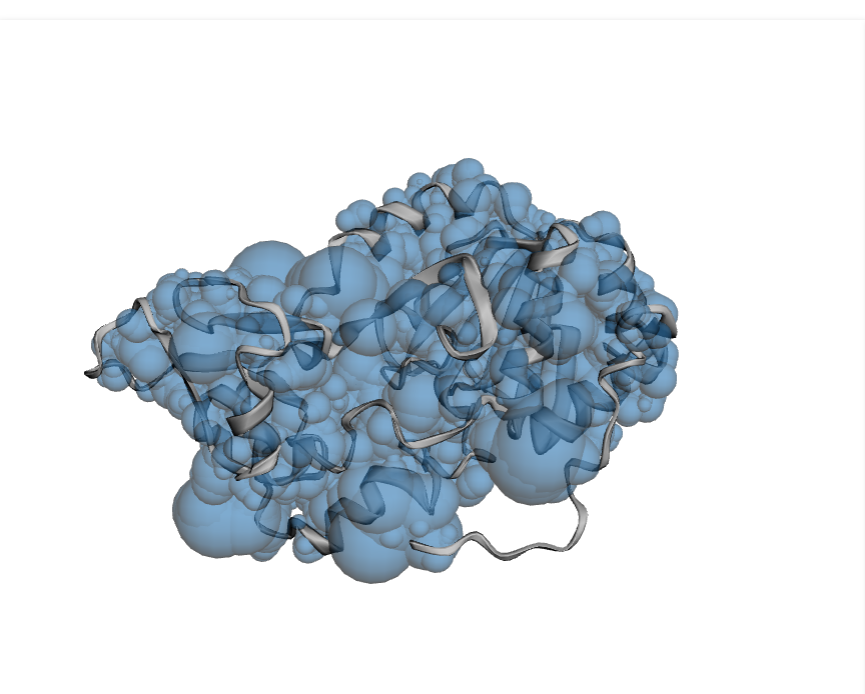

**Supplementary Figure 3:** Active site of SIRT1 receptor; Sky blue spheres represent the active site region.

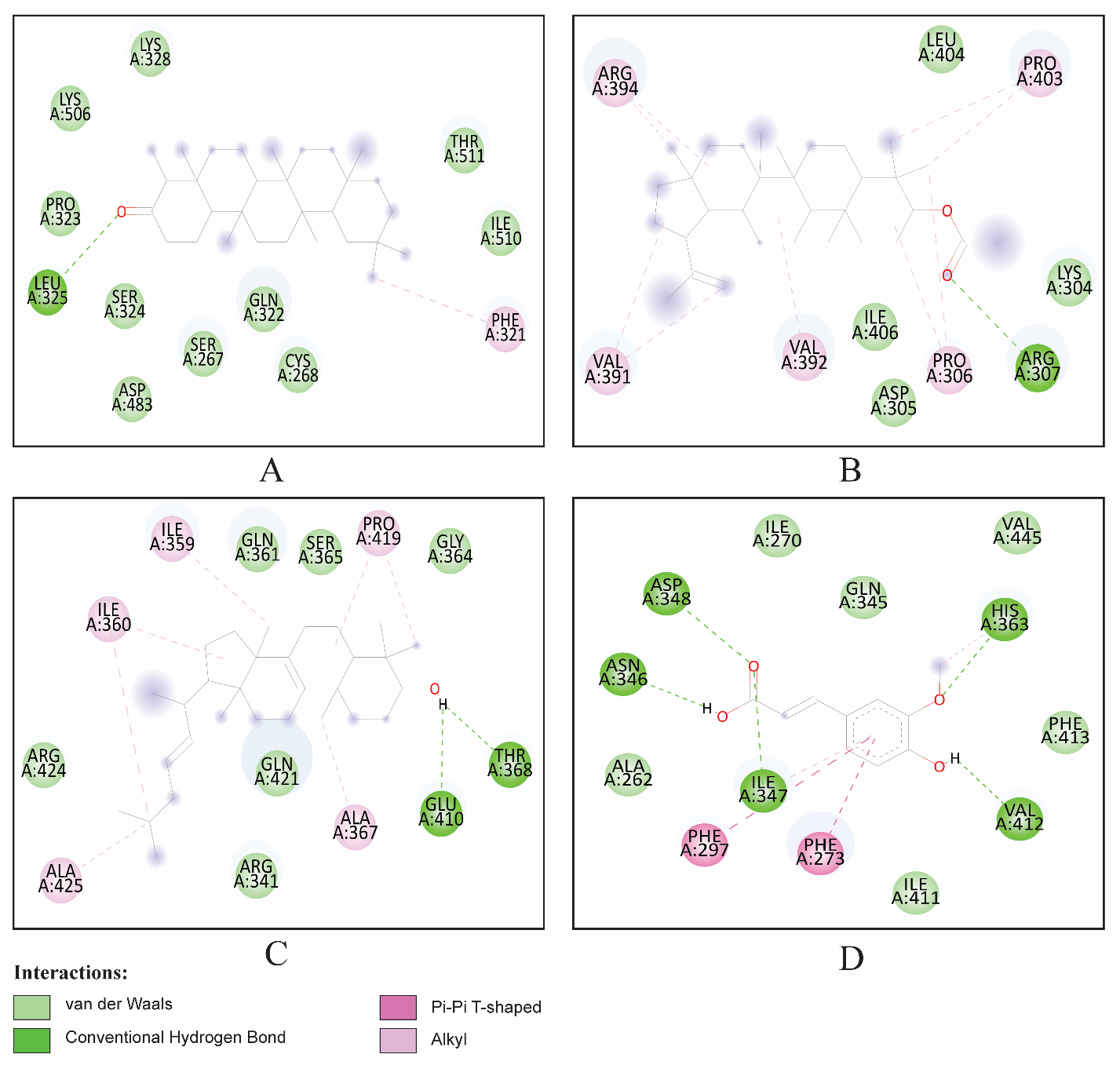

**Supplementary Figure 4:** 2D interaction between the protein ligand complex. Here, (A) Friedelin, (B) Lupeol acetate, (C) Gluanol, (D) Ferulic acid, figure showing the ligand contact with the protein SIRT1 after molecular docking.

**Supplementary Table 1: Chemical properties of the SIRT1 compounds.**

| **Compound Name** | **PubChem CID** | **Molecular Formula** | **Canonical SMILES** | **IUPAC NAME** | **Reference** |
| --- | --- | --- | --- | --- | --- |
| Ascorbic acid | 54670067 | C6H8O6 or HC6H7O6 | C(C(C1C(=C(C(=O)O1)O)O)O)O | (2R)-2-[(1S)-1,2-dihydroxyethyl]-3,4-dihydroxy-2H-furan-5-one | <https://www.sciencedirect.com/science/article/pii/S2667142522000975#bib0034> |
| B-sitosterol | 222284 | C29H50O | CCC(CCC(C)C1CCC2C1(CCC3C2CC=C4C3(CCC(C4)O)C)C)C(C)C | (3S,8S,9S,10R,13R,14S,17R)-17-[(2R,5R)-5-ethyl-6-methylheptan-2-yl]-10,13-dimethyl-2,3,4,7,8,9,11,12,14,15,16,17-dodecahydro-1H-cyclopenta[a]phenanthren-3-ol | <http://nopr.niscpr.res.in/bitstream/123456789/3777/1/NPR%208%281%29%2084-90.pdf> |
| Catechin | 9064 | C15H14O6 | C1C(C(OC2=CC(=CC(=C21)O)O)C3=CC(=C(C=C3)O)O)O | (2R,3S)-2-(3,4-dihydroxyphenyl)-3,4-dihydro-2H-chromene-3,5,7-triol | [http://nopr.niscpr.res.in/bitstream/123456789/3777/1/NPR%208%281%29%2084-90](http://nopr.niscpr.res.in/bitstream/123456789/3777/1/NPR%208%281%29%2084-90.pdf) |
| Chlorogenic acid | 1794427 | C16H18O9 | C1C(C(C(CC1(C(=O)O)O)OC(=O)C=CC2=CC(=C(C=C2)O)O)O)O | (1S,3R,4R,5R)-3-[(E)-3-(3,4-dihydroxyphenyl)prop-2-enoyl]oxy-1,4,5-trihydroxycyclohexane-1-carboxylic acid | <http://nopr.niscpr.res.in/bitstream/123456789/3777/1/NPR%208%281%29%2084-90.pdf> |
| Ferulic acid | 445858 | C10H10O 3704 | COC1=C(C=CC(=C1)C=CC(=O)O)O | (E)-3-(4-hydroxy-3-methoxyphenyl)prop-2-enoic acid | <http://nopr.niscpr.res.in/bitstream/123456789/3777/1/NPR%208%281%29%2084-90.pdf> |
| Friedelin | 91472 | C30H50O | CC1C(=O)CCC2C1(CCC3C2(CCC4(C3(CCC5(C4CC(CC5)(C)C)C)C)C)C)C | (4R,4aS,6aS,6aS,6bR,8aR,12aR,14aS,14bS)-4,4a,6a,6b,8a,11,11,14a-octamethyl-2,4,5,6,6a,7,8,9,10,12,12a,13,14,14b-tetradecahydro-1H-picen-3-one | <http://nopr.niscpr.res.in/bitstream/123456789/3777/1/NPR%208%281%29%2084-90.pdf> |
| Gallic acid | 370 | C7H6O5 | C1=C(C=C(C(=C1O)O)O)C(=O)O | 3,4,5-trihydroxybenzoic acid |  |
| Gluanol | 101316952 | C30H50O | CC(C)CC=CC(C)C1CCC2(C1(CCC3=C2CCC4C3(CCC(C4(C)C)O)C)C)C | (3S,5R,10S,13S,14S,17S)-4,4,10,13,14-pentamethyl-17-[(E,2S)-6-methylhept-3-en-2-yl]-2,3,5,6,7,11,12,15,16,17-decahydro-1H-cyclopenta[a]phenanthren-3-ol | <https://doi.org/10.1016/j.crgsc.2020.100020> |
| Hentriacontane | 12410 | C31H64 | CCCCCCCCCCCCCCCCCCCCCCCCCCCCCCC | hentriacontane | https://doi.org/10.1016/j.crgsc.2020.100021 |
| Lupeol acetate | 92157 | C32H52O2 | CC(=C)C1CCC2(C1C3CCC4C5(CCC(C(C5CCC4(C3(CC2)C)C)(C)C)OC(=O)C)C)C | [(1R,3aR,5aR,5bR,7aR,9S,11aR,11bR,13aR,13bR)-3a,5a,5b,8,8,11a-hexamethyl-1-prop-1-en-2-yl-1,2,3,4,5,6,7,7a,9,10,11,11b,12,13,13a,13b-hexadecahydrocyclopenta[a]chrysen-9-yl] acetate | https://doi.org/10.1016/j.crgsc.2020.100022 |
| Quercetin | 5280459 | C21H20O11 | C1=CC(=C(C=C1C2=C(C(=O)C3=C(C=C(C=C3O2)O)O)O)O)O | 2-(3,4-dihydroxyphenyl)-3,5,7-trihydroxychromen-4-one | https://doi.org/10.1016/j.crgsc.2020.100024 |
| Taraxasterol | 115250 | C30H50O | CC1C2C3CCC4C5(CCC(C(C5CCC4(C3(CCC2(CCC1=C)C)C)C)(C)C)O)C | (3S,4aR,6aR,6aR,6bR,8aR,12S,12aR,14aR,14bR)-4,4,6a,6b,8a,12,14b-heptamethyl-11-methylidene-1,2,3,4a,5,6,6a,7,8,9,10,12,12a,13,14,14a-hexadecahydropicen-3-ol | https://doi.org/10.1016/j.crgsc.2020.100025 |
| Tiglic acid | 125468 | C5H8O2 | CC=C(C)C(=O)O | (E)-2-methylbut-2-enoic acid | https://doi.org/10.1016/j.crgsc.2020.100026 |

**Supplementary Table 2: ADMET properties of the compounds.**

| **ADMET** | **Properties** | **Ascorbic acid** | **B-sitosterol** | **Catechin** | **Chlorogenic acid** | **Ferulic acid** | **Friedelin** | **Gallic acid** | **Gluanol** | **Lupeol acetate** | **Quercetin** | **Taraxasterol** | **Tiglic acid** |
| --- | --- | --- | --- | --- | --- | --- | --- | --- | --- | --- | --- | --- | --- |
| Adsorption | Human intestinal absorption | 0.069 | 0.004 | 0.096 | 0.873 | 0.03 | 0.157 | 0.085 | 0.01 | 0.009 | 0.014 | 0.008 | 0.009 |
|  | Caco-2 permeability | -5.917 | -4.756 | -5.971 | -6.127 | -4.902 | -5.159 | -5.728 | -4.783 | -4.949 | -5.204 | -5.028 | -4.588 |
|  | MDCK | 0.000143 | 8.63E-06 | 4.27E-06 | 9.88E-05 | 1.46E-05 | 5.88E-06 | 5.11E-06 | 8.07E-06 | 8.57E-06 | 7.69E-06 | 8.08E-06 | 2.67E-05 |
|  | F20% | 0.918 | 0.01 | 0.99 | 0.822 | 0.047 | 0.712 | 0.964 | 0.006 | 0.037 | 0.93 | 0.38 | 0.003 |
| Distribution | Blood brain Permeability | -0.985 | 0.781 | -1.054 | -1.407 | -0.239 | 0.72 | -1.102 | 0.711 | 0.644 | -1.407 | 0.705 | -0.248 |
|  | Fraction unbound in plasma | 0.825 | 0 | 0.235 | 0.658 | 0.343 | 0 | 0.617 | 0 | 0 | 0.658 | 0 | 0.692 |
|  | Volume of distribution (Vd) (L/kg) | 0.218 | 0.193 | 1.027 | 0.581 | -1.367 | -0.272 | -1.855 | 0.58 | -0.12 | 0.581 | -0.048 | -0.763 |
|  | CNS permeability | -3.217 | -1.705 | -3.298 | -3.856 | -2.612 | -1.555 | -3.74 | -1.91 | -1.754 | -3.856 | -1.668 | -2.53 |
| Metabolism | CYP1A2-inh | 0.013 | 0.044 | 0.219 | 0.036 | 0.059 | 0.02 | 0.023 | 0.026 | 0.021 | 0.943 | 0.029 | 0.046 |
|  | CYP1A2-sub | 0.052 | 0.491 | 0.295 | 0.042 | 0.478 | 0.63 | 0.075 | 0.521 | 0.406 | 0.115 | 0.49 | 0.471 |
|  | CYP2C9-inh | 0.007 | 0.096 | 0.218 | 0.016 | 0.142 | 0.061 | 0.188 | 0.117 | 0.07 | 0.598 | 0.08 | 0.054 |
|  | CYP2C9-sub | 0.23 | 0.314 | 0.838 | 0.511 | 0.367 | 0.39 | 0.061 | 0.736 | 0.392 | 0.643 | 0.236 | 0.317 |
| Excretion | CL | 9.964 | 16.686 | 17.911 | 3.251 | 7.48 | 19.969 | 10.108 | 14.915 | 7.779 | 8.284 | 16.329 | 5.154 |
|  | T 1/2 | 0.928 | 0.013 | 0.853 | 0.928 | 0.926 | 0.033 | 0.947 | 0.007 | 0.009 | 0.929 | 0.01 | 0.897 |
| Toxicity | Hepatotoxicity | Inactive | Inactive | Inactive | Inactive | Inactive | Inactive | Inactive | Inactive | Inactive | Inactive | Inactive | Inactive |
|  | Carcinogenicity | Inactive | Inactive | Inactive | Inactive | Inactive | Inactive | **Active** | Inactive | Inactive | **Active** | Inactive | Inactive |
|  | Mutagenicity | Inactive | Inactive | Inactive | Inactive | Inactive | Inactive | Inactive | Inactive | Inactive | **Active** | Inactive | Inactive |
|  | Cytotoxicity | Inactive | Inactive | Inactive | Inactive | Inactive | Inactive | Inactive | Inactive | Inactive | Inactive | Inactive | Inactive |
